# CRK5 preserves antioxidant homeostasis and prevents cell death during dark-induced senescence through inhibiting the salicylic acid signaling pathway

**DOI:** 10.64898/2026.01.12.698963

**Authors:** Muhammad Kamran, Paweł Burdiak, Anna Rusaczonek, Roshanak Zarrin Ghalami, Stanisław M. Karpiński

## Abstract

**Background:** Dark-induced senescence (DIS) is a widely used model for dissecting the regulatory mechanisms that manage leaf aging, redox imbalance, and cell death (CD) in plants. Salicylic acid (SA) is a central hormonal regulator of these processes. However, the mechanism involving upstream components, which integrate SA-dependent pathways with antioxidant homeostasis during DIS, remains unresolved. CYSTEINE-RICH RECEPTOR-LIKE KINASE 5 (CRK5) is a membrane-localized protein that plays a role in developmental and stress-responsive pathways. Its promoter contains multiple W-box *cis*-elements, indicating regulation by WRKY factors in SA-mediated pathways. This study investigates how CRK5 modulates SA-dependent CD and antioxidant dynamics during DIS.

**Results:** In this study, SA-accumulating mutant *crk5* exhibited accelerated senescence, elevated electrolyte leakage, enhanced micro-lesion formation, and markedly increased reactive oxygen species (ROS) accumulation under both control and dark conditions. These phenotypes were accompanied by a substantial reduction in carotenoid and xanthophyll pools, enhanced accumulation of phenolic compounds, and increased free radical scavenging capacity, including ascorbate peroxidase, catalase, and superoxide dismutase activities. Importantly, *crk5* phenotype was fully reverted in *crk5sid2* and *crk5*NahG double mutants, confirming that *crk5* DIS phenotype is induced by activation of the SA-signaling pathway.

Transcriptome profiling revealed extensive deregulation of senescence-, CD-, and redox-associated genes in *crk5* during darkness, including strong induction of SAGs, metacaspases, autophagy, and antioxidant-related transcripts. The line with constitutively enhanced SA level (*cpr1*), used as a control, showed similar phenotypes to *crk5*, although transcriptional reprogramming was largely absent in *cpr1* after darkness, highlighting CRK5 as a key upstream negative regulator of SA-mediated CD and positive regulator of antioxidant homeostasis.

**Conclusion:** Our work presents CRK5 as a central regulatory hub that inhibits the SA-signaling, ROS burst, and CD activation during DIS. Loss of CRK5 function is associated with the activation of SA-signaling, altered antioxidant systems, increased ROS burden, and ROS-driven CD acceleration, resulting in accelerated senescence. Conversely, suppression of SA-biosynthesis or -catabolism in a *crk5* background restores the wild-type phenotype. These findings position this receptor kinase as a key mediator that coordinates hormonal, metabolic, and oxidative pathways to maintain leaf viability, providing mechanistic insight into the control of stress-induced senescence and CD in Arabidopsis.

## Background

Leaf senescence represents the final, essential stage of foliar development, characterized by a fundamental functional transition from active nutrient assimilation to systematic nutrient remobilization, which is critical for overall plant fitness and survival. This process is highly regulated, following a predetermined program of events often classified as cell death (CD) [1, 2]. In plant biology, dark-induced senescence (DIS) is frequently utilized as a critical experimental model system to study natural age-related senescence. This artificial induction effectively promotes many typical senescence symptoms, including extensive chlorophyll degradation, altered photosystem II performance, and organized protein catabolism [2, 3].

Understanding the manner in which DIS triggers the controlled collapse of antioxidant defenses is vital to defining the regulatory frontiers governing plant longevity [4]. Reactive Oxygen Species (ROS) are unavoidable byproducts of aerobic metabolism [5]. They include highly reactive molecules such as hydrogen peroxide (H_2_O_2_), superoxide (O^2−^), and the hydroxyl radical (OH^−^). In the context of senescence, ROS exhibit a dual function. At low, controlled concentrations, they act as indispensable signaling molecules, actively initiating the senescence program, whereas at high levels they trigger oxidative damage to nucleic acids, lipids, and proteins, leading to irreversible injury and CD [5]. Consequently, leaves undergoing senescence are typically characterized by an acute imbalance: excessive ROS accumulation coupled with a significant reduction in the cellular capacity for antioxidant scavenging [6]. Maintaining cellular redox homeostasis relies on balancing ROS production with immediate scavenging by antioxidant systems [7]. When scavenging systems become overwhelmed, the delicate redox balance is permanently disturbed, pushing the cell toward self-destruction [8]. Plants under stress conditions significantly aggravate the generation of ROS, creating a persistent challenge to cellular redox homeostasis that ultimately dictates the transition from stress tolerance to CD [9].

CD is an active, conserved mechanism of cellular suicide that is central to both normal plant development and environmental stress management, commonly preceded by morphological changes such as nuclear condensation [10]. The activation of the CD process requires the creation of a specific, oxidizing redox environment in various cellular compartments. Research has focused on understanding the interplay between ROS and the complex network of antioxidant molecules and enzymes necessary to maintain the redox environment required for cell survival [11]. Studies on Arabidopsis have demonstrated that H_2_O_2_ directly induces CD, confirming the central role of antioxidant capacity in determining whether ROS serves as an adaptive signal or a death trigger [12].

This critical life-or-death decision, which depends on chloroplasts and mitochondria membrane stability, is regulated by the efficiency and coordination of both enzymatic and non-enzymatic defenses [13–15]. Key enzymatic antioxidant, such as Superoxide Dismutase (SOD), serves as the primary defensive line by converting highly reactive superoxide (O^2−^) into the more stable signaling molecule, hydrogen peroxide (H_2_O_2_) [12]. Downstream H_2_O_2_ scavengers, including Catalase (CAT) and Ascorbate Peroxidase (APX), function to maintain H_2_O_2_ at basal signaling concentrations, essential for acclimation [9, 16]. Concurrently, non-enzymatic antioxidants, encompassing potent radical scavengers such as phenolic compounds and critical photo-protective agents like carotenoids, establish a vital constitutive defense buffer [17]. Two main classes of carotenoids are carotene and xanthophyll, which have been described in their roles as oxidative protectants [18, 19]. Genetic evidence from Arabidopsis, notably the *radical-induced cell death1* (*rcd1*) mutant, reveals that the constitutive upregulation and enhanced redox cycling of phenolic compounds confer remarkable resistance to oxidative stress and subsequent CD, thereby actively blocking the CD cascade [20, 21]. The regulatory role of signaling molecules, particularly salicylic acid (SA), in modulating the timing of the shift in redox balance is crucial in determining cell fate during DIS. The central question addressed by detailed mechanistic studies is how DIS specifically manipulates the antioxidant machinery to manage this shift, from maintaining a highly regulated, low-level signaling ROS pool to permitting the accumulation of damaging ROS necessary for the final execution phase of CD [22, 23].

Salicylic acid (SA) serves as a pivotal phytohormone, coordinating numerous physiological and biochemical processes throughout the entire plant’s lifespan. Its regulatory influence spans vegetative growth, photosynthesis, respiration, thermogenesis, and, critically, both senescence and a specific type of CD that is independent of the canonical hypersensitive response [24–26]. An important aspect of SA’s activity is its contribution to maintaining cellular redox homeostasis, primarily achieved through the modulation of antioxidant enzyme activity [27]. Studies investigating SA’s influence on plant physiology, such as those conducted during acclimation to high light stress, reveal a tight regulatory relationship between SA levels, H_2_O_2_ content, and major non-enzymatic antioxidant pools. Specifically, high endogenous foliar SA concentrations in Arabidopsis mutants are found to correlate strictly with elevated levels of H_2_O_2_. Crucially, this elevation in the oxidative signal is paralleled by a higher content of reduced glutathione (GSH) [28]. This synchronized increase in both the primary H_2_O_2_ and the major non-enzymatic protective buffer (GSH) suggests that SA is not merely inducing stress but is actively coordinating a state of high-flux redox signaling. By increasing the capacity of GSH pool, which is essential for recycling ascorbic acid (AsA) and supporting H_2_O_2_ detoxification, the H_2_O_2_ signal is received without causing immediate, widespread catastrophic oxidative failure [28, 29].

It has been shown that mutants with lower SA levels exhibit better fitness, produce more seeds, and accumulate more biomass, whereas mutants with higher SA content often exhibit a dwarf phenotype [30–33]. Interestingly, some mutants with decreased SA biosynthesis are more susceptible to biotrophic pathogen infection (i.e., *eds5* and *sid2*), while others are more resistant to abiotic stresses (i.e., *eds1* or *pad4*) [30, 34, 35]. In response to stress, the *lsd1* mutant exhibits elevated SA-accumulation and increased CD due to activation of EDS1 and PAD4, because these phenotypes are suppressed in the *lsd1/eds1* and *lsd1/pad4* double mutants, which also show markedly reduced SA and ROS levels [30, 36–39]. These findings suggest that the role of SA in response to biotic and abiotic stresses differs, and it has been postulated that SA acts as a double-edged sword for CD in plants [40]. SA-biosynthesis and -signaling pathways are tightly regulated and integrate multiple upstream signals, including those perceived by receptor-like kinases (RLKs) at the cell surface [41, 42].

Cysteine-rich Receptor-Like Kinases (CRKs) constitute a distinctive subfamily of Receptor-Like Kinases (RLKs) that contain multiple cysteine residues in the extracellular domain (DUF26) organized in conserved motifs [43]. The structural configuration of CRKs provides a powerful conceptual framework for their role in environmental sensing. Due to the high abundance and reactivity of nucleophilic thiol groups within the cysteine residues, CRKs are considered excellent candidates for functioning as extracellular ROS sensors, regulating signal transduction pathways related to plant defense and acclimation, stomatal closure, CD, and developmental senescence [42, 44, 45]. By directly linking extracellular oxidative signals to intracellular kinase activity, CRKs serve as fundamental convergence points for converting stress-induced redox perturbations into regulated defense and survival responses, thereby directly influencing the maintenance of cellular redox homeostasis and the status of the antioxidant system [42]. CYSTEINE-RICH RECEPTOR-LIKE KINASE 5 (CRK5) in Arabidopsis has established its critical role in regulating growth, senescence, thermoregulation, and acclimatory responses under basal and abiotic stress conditions [43, 46]. CRK5 plays an essential function in plant defense against biotic stimuli, specifically by regulating the SA-signaling pathway, via the MPK3/6–WRKY70–TGA2/6 cascade, positioning CRK5 upstream of the MPK module in response to *Vd*-toxins [47].

The objective of this study was to check the relationship between SA-mediated antioxidant homeostasis during CD through CRK5-mediated negative regulation of dark-induced senescence. We hypothesized that during dark-induced senescence, *crk5* plants would exhibit enhanced CD and more pronounced alterations in antioxidant activity, driven by their elevated SA-accumulation relative to control conditions. To prove this, we used single and double mutant lines impaired in CRK5 activity and deficient in the SA-synthesis or -catabolism (*sid2* and NahG). The *cpr1* (constitutive expresser of pathogenesis-related genes 1) mutant was used as a control, since it has increased levels of SA and elevated expression of pathogenesis-related (PR) genes. The CPR1 is a part of the ubiquitin ligase complex and acts upstream of SA, a negative regulator of plant defense and immune responses [48, 49]. Moreover, our data showed a higher CD rate, reduced pigment content, altered enzymatic and non-enzymatic antioxidant activity, and higher ROS accumulation in SA-accumulating lines (*crk5* and *cpr1*) compared to the wild-type. The RNA-seq data supported our hypothesis regarding the proposed inhibitory role of CRK5 in these processes. Taken together, our data indicate that CRK5 acts as a negative regulator of the SA-signaling pathway, inhibiting accelerated DIS and CD, and altering the antioxidant and redox homeostasis.

## Materials and methods

### Plant Material and Growing Conditions

All Arabidopsis lines used in this study were in the Columbia-0 (Col-0) genetic background. Double mutant lines were generated through genetic crosses, and primers employed for genotyping are listed in Table S1. Seeds were stratified at 4 °C for three days prior to germination. Plants were then cultivated in a controlled growth chamber on Jiffy Pots under long-day conditions (16 h light/8 h dark) with a constant light intensity of 120 µmol photons m⁻² s⁻¹, at 22 °C and 70 ± 5% relative humidity, for 4–5 weeks. To induce dark-associated senescence, four-week-old plants were transferred to continuous darkness for a period of four days.

### Relative electrolyte leakage

Leaves were detached and placed into 50 mL Falcon tubes containing 35 mL of Milli-Q water (Merck Millipore, Darmstadt, Germany). Electrolyte leakage was determined using a conductivity meter (WTW INOLAB Cond Level 1, Weilheim, Germany). Relative electrolyte leakage was calculated as the ratio of conductivity measured after 1 h of incubation to the total leakage obtained following autoclaving of the samples.

### Trypan blue staining

A trypan blue (TB) stock solution (30 µmol; Sigma-Aldrich, St. Louis, MO, USA) was prepared in a mixture of lactic acid, glycerol, and water (10:10:20, v/v/v) and diluted 1:2 with 96% ethanol to obtain the working solution. Leaves from control and dark-treated plants were harvested and immediately immersed in the TB working solution in 50 mL Falcon tubes. Samples were incubated for 30 min at room temperature with gentle agitation. Following staining, the TB solution was discarded and replaced with methanol. Leaves were destained in methanol for 24 h, with the methanol refreshed several times to ensure complete removal of chlorophyll. Chlorophyll-depleted leaves were imaged using a Nikon SMZ18 stereomicroscope equipped with a Nikon d5100 camera (Nikon Inc., Melville, NY, USA). Images were analyzed using ImageJ software (version 1.8.0), and micro-lesions (trypan blue–positive spots) were quantified per mm² of leaf area.

### Pigment content analysis

Frozen leaf material (20–50 mg) was ground using a Mixer Mill MM 400 (Retsch, Düsseldorf, Germany) for 5 min at 30 Hz and 4 °C in the presence of 1 mL pre-chilled acetone (−20 °C). The resulting extract was evaporated to dryness using a Savant DNA120 SpeedVac concentrator (Thermo Scientific, Waltham, MA, USA). The residue was subsequently resuspended in cold solvent A (acetonitrile:methanol, 90:10, v/v) and briefly homogenized for 1 min. After filtration through a 0.2 μm nylon membrane filter (Whatman, Marlborough, MA, USA), samples were transferred to autosampler vials, sealed, and stored at −80 °C in darkness until HPLC analysis.

Pigment separation was performed using a Shimadzu HPLC system (Kyoto, Japan) equipped with a Synergi™ MAX-RP column (4 μm, 80 Å, 250 × 4.6 mm; Phenomenex, Torrance, CA, USA) maintained at 30 °C. Xanthophylls were eluted using solvent A for 10 min, followed by elution of carotenes with solvent B (methanol:ethyl acetate, 68:32, v/v) for an additional 10 min at a constant flow rate of 1 mL min⁻¹. Individual pigments were identified by comparison with authentic standards (Sigma-Aldrich, St. Louis, MO, USA; Hoffmann-La Roche, Basel, Switzerland; Fluka, Buchs, Switzerland). Total carotenoid content was calculated as the sum of quantified xanthophylls and carotenes and expressed as peak area per μg of DW, following previously established protocols [50, 51].

### ABTS and DPPH scavenging activity

The antioxidant capacity of methanol extracts was assessed by evaluating their ability to scavenge ABTS and DPPH free radicals, as well as their ferric reducing power using the FRAP assay. Freeze-dried tissue was finely powdered, extracted with methanol, and centrifuged at 13,000 rpm for 15 min at room temperature. The resulting supernatant was collected and used for antioxidant activity measurements.

ABTS radical scavenging activity was determined according to the method originally described by [52]. The ABTS•⁺ working solution was generated by incubating an aqueous mixture of 7 mM ABTS (2,2′-azinobis-(3-ethylbenzothiazoline-6-sulfonic acid)) and 2.45 mM potassium persulfate in darkness at room temperature for 16 h. Prior to analysis, the solution was diluted with ethanol to obtain an absorbance of 0.70 ± 0.02 at 734 nm. Sample extracts were mixed with the ABTS•⁺ solution, and absorbance was recorded at 734 nm after 6 min using a microplate reader (Multiskan GO, Thermo Scientific, Waltham, MA, USA).

DPPH radical scavenging activity was measured following the established procedure of [53]. The extracts of the prepared samples were incubated with an ethanol solution of DPPH• (2,2-diphenyl-1-picrylhydrazyl), and absorbance was measured at 517 nm after 20 min of reaction. Radical scavenging capacity was calculated as the percentage inhibition relative to the control.

Antioxidant activity for both ABTS and DPPH assays was calculated using the equation: (%) inhibition = [(A₀ − A₁)/A₀] × 100, where A₀ and A₁ represent the absorbance of the control and sample, respectively. Results were expressed as μmol Trolox equivalents (TE) per gram of DW.

Ferric reducing antioxidant power (FRAP) was determined by measuring the reduction of Fe³⁺–TPTZ to Fe²⁺–TPTZ according to the method described by [52].

### Phenolic Profile

Total phenolic compounds were quantified following the extraction of 100 mg of plant tissue with 80% (v/v) methanol. Phenolic content was determined using the Folin–Ciocalteu colorimetric assay, according to [54]. After centrifugation, aliquots of the extracts were combined with deionized water, Folin–Ciocalteu reagent, and saturated sodium carbonate (Na₂CO₃). The reaction mixtures were incubated at 40 °C for 30 min, cooled to room temperature, and the absorbance was measured at 740 nm. Total polyphenol content was expressed as milligrams of gallic acid equivalents (GAE) per 100 g DW.

Phenylpropanoids, flavonols, and anthocyanins were quantified using the spectrophotometric method described by [55]. Methanol extracts were mixed with acidified ethanol (0.1% HCl in 96% ethanol) and aqueous HCl (2% HCl in water). After a 15 min incubation, absorbance was recorded at 320 nm, 360 nm, and 520 nm using a BioSpectrometer kinetic system. Caffeic acid, quercetin, and cyanidin were used as calibration standards for phenylpropanoids, flavonols, and anthocyanins, respectively. The concentrations of individual phenolic groups were expressed as milligrams per 100 g DW.

### Hydrogen Peroxide Levels Determination

Hydrogen peroxide (H₂O₂) content was quantified following the method of [56] with minor modifications. Approximately 50 mg of leaf tissue was homogenized in a TissueLyser LT (Qiagen, The Netherlands) for 5 min at 50 Hz and 4 °C in 300 µL of ice-cold 0.1% (w/v) trichloroacetic acid. The homogenate was centrifuged at 13,000 rpm for 15 min, and the resulting supernatant was collected for analysis. An aliquot of the extract was combined with 10 mM potassium phosphate buffer (pH 7.0) and 1 M potassium iodide (KI) in a volumetric ratio of 1:1:2. Absorbance was recorded at 390 nm using a Multiscan GO microplate reader (Thermo Scientific, USA). Hydrogen peroxide concentration was calculated from a standard calibration curve and expressed as µmol H₂O₂ per 100 mg DW.

Histochemical detection of H₂O₂ was performed using 3,3′-diaminobenzidine (DAB) staining as described by [57]. Excised leaves were immersed in a staining solution containing 0.1% (w/v) DAB, 2 mM dimethyl sulfoxide (DMSO), and 0.05% (v/v) Tween-20 prepared in Milli-Q water (pH 3.8) and incubated for 4 h at room temperature. Following staining, chlorophyll was removed by incubation in 0.25% (w/v) chloral hydrate for 24 h. Stained leaves were subsequently examined using a Leica M165 FC stereomicroscope.

### Protein extraction

Leaf samples were detached, immediately frozen in liquid nitrogen, and 50–100 mg portions homogenized using a TissueLyser LT (Qiagen) at 50 s⁻¹ for 5 min at 4°C in 1 ml extraction buffer (100 mM Tricine-Tris, pH 7.5, 3 mM MgSO₄, 3 mM EGTA, 1 mM DTT). Samples were incubated on ice for 15 min, then centrifuged at 18,000 × g for 20 min. The resulting supernatant served as the enzyme extract and was stored at −80°C for subsequent assays. For ascorbate peroxidase activity measurements, the extract was supplemented with 5 mM L-ascorbate. Protein concentrations were quantified via the Bradford (1976) method [58] using a commercial protein assay kit (Fermentas) standardized against bovine serum albumin (Fermentas).

### Determination of antioxidant enzyme activities

APX activity was determined following [59] with modifications. Enzyme extract was combined with assay buffer (50 mM phosphate, pH 7.0, 1 mM EDTA, 15 mM L-ascorbate, 40 mM H₂O₂) at ratios ensuring 20–80% absorbance decline. Activity was calculated from the ascorbate oxidation rate over 2 min, using its extinction coefficient at 290 nm (ε =2.8 mM^−1^cm^−1^). Values are reported as μmol L-AsA mg⁻¹ protein min⁻¹.

CAT activity was performed according to [60] with modifications. A 30% (v/v) H₂O₂ solution was diluted in 50 mM phosphate buffer (pH 7.0) to an absorbance of 0.5 (±0.02) at 240 nm (initial concentration of H_2_O_2_ ca. 13 mM). Enzyme extract was added at ratios yielding 20–80% absorbance decrease. Activity was measured by H₂O₂ decomposition rate over 2 min, using its extinction coefficient of H_2_O_2_ at 240 nm (ε =43.6 mM^−1^cm^−1^). Results are expressed as μmol H₂O₂ decomposed mg⁻¹ protein min⁻¹.

SOD activity was assayed spectrophotometrically per [61]. Freshly prepared enzyme assay mixture contained 0.1 M phosphate buffer (pH 7.5), 2.4 μM riboflavin, 840 μM nitroblue tetrazolium (NBT) salt, 150 mM methionine, 12 mM Na_2_EDTA in the proportion 8:1:1:1:1 by vol., respectively. Enzyme extract was added to achieve 20–80% inhibition of NBT reduction. After 15 min illumination (500 μmol m⁻² s⁻¹ white LED; Photon System Instruments SL 3500-W-D) or dark incubation (blank), absorbance was measured at 560 nm. Results were expressed in units (U) defined as the amount of enzyme that inhibits NBT photo-reduction to blue formazan by 50% *per* 1 mg of protein.

All assays were performed using a UV-Vis Multiskan GO Microplate Spectrophotometer (Thermo Scientific, Waltham, MA, USA).

### Library preparation and RNA-seq analysis

For RNA isolation, 4- or 4.5-week-old plants were used. The experiment was performed on three biological replicates. Each replicate consisted of several pooled detached leaves from at least three individual plants. Frozen leaves were ground using a mortar and pestle in liquid nitrogen, and RNA was isolated using the Spectrum™ Plant Total RNA Kit (Sigma, St. Louis, MO, USA). mRNA fraction was isolated using Oligo d(T)25 Magnetic Beads (NEB, S1419S) from 20 ug of total RNA. RNA-seq libraries were self-prepared according to [62]. RNA-seq libraries were sequenced in 150-bp paired-end reads using the Aviti High flowcell platform (Element Biosciences), conducted by Genomed (Warsaw, Poland). Unique Molecular Identifiers (UMIs) were extracted from R2 reads using UMI-tools extract [63]. Subsequently, R1 reads underwent adapter trimming and quality filtering to remove low-quality sequences using Trim Galore (https://github.com/FelixKrueger/TrimGalore). The cleaned reads were then aligned using STAR to the *Arabidopsis thaliana* reference genome (Ensembl Plants release 60) [64]. To eliminate PCR duplicates, UMI-tools dedup was employed, leveraging both UMI sequences and mapping positions. Gene-level read counts were generated using htseq-count [65]. The resulting count data were imported into R, where a count matrix was constructed. Differential expression analysis was performed using the DESeq2 package [66]. Significantly enriched GO terms were identified in up-/downregulated gene sets using the package (v4.10.1) [67].

### Statistical analysis

The statistical analysis for ion leakage, micro-lesions, pigment content, scavenging activities (DPPH and ABTS), Phenolic content, H_2_O_2_ content, and antioxidant activities (APX, CAT, and SOD) was performed using GraphPad Prism 8 software.

### Accession numbers

Accession numbers of all genes in Fig. 5C and D are given in Table S2.

## Results

### CRK5, through the prevention of SA-signaling, accelerates dark-induced senescence and cell death

To clarify the role of salicylic acid signaling in CRK5-mediated inhibition of dark-induced senescence and cell death, we analyzed mutants with elevated SA levels (*crk5* and *cpr1*), SA-synthesis or -catabolism deficient lines (*sid2* and NahG), and the corresponding double mutants (*crk5sid2* and *crk5*NahG) (Fig. S1). We observed induced accelerated senescence in plants with elevated SA content under dark conditions (Fig. 1A). Subsequently, we measured electrolyte leakage to assess CD in these genotypes. We found higher ion leakage levels in mutants with elevated SA content (*crk5* and *cpr1*) even in control conditions, compared to the wild-type (Col-0). In contrast, the CD level was not significantly different in SA-synthesis or -catabolism deficient mutants (*sid2* and NahG) and double mutants (*crk5sid2* and *crk5*NahG). In the dark-treated plants, these changes were even more pronounced in the *crk5* and *cpr1* mutants. (Fig. 1B). Moreover, micro-lesion formation was assessed using TB staining. Micro-lesions constitute small lesion areas within the leaf tissue, comprising one or a couple of dead cells [68, 69]. In control conditions, micro-lesions were consistently higher in *crk5* and *cpr1*. Under dark-induced senescence, these differences became even more pronounced compared with Col-0 and SA-synthesis or -catabolism deficient lines (Fig. 1C, D). These results indicate that SA-signaling during DIS enhances senescence and promote CD.

**Figure 1.**
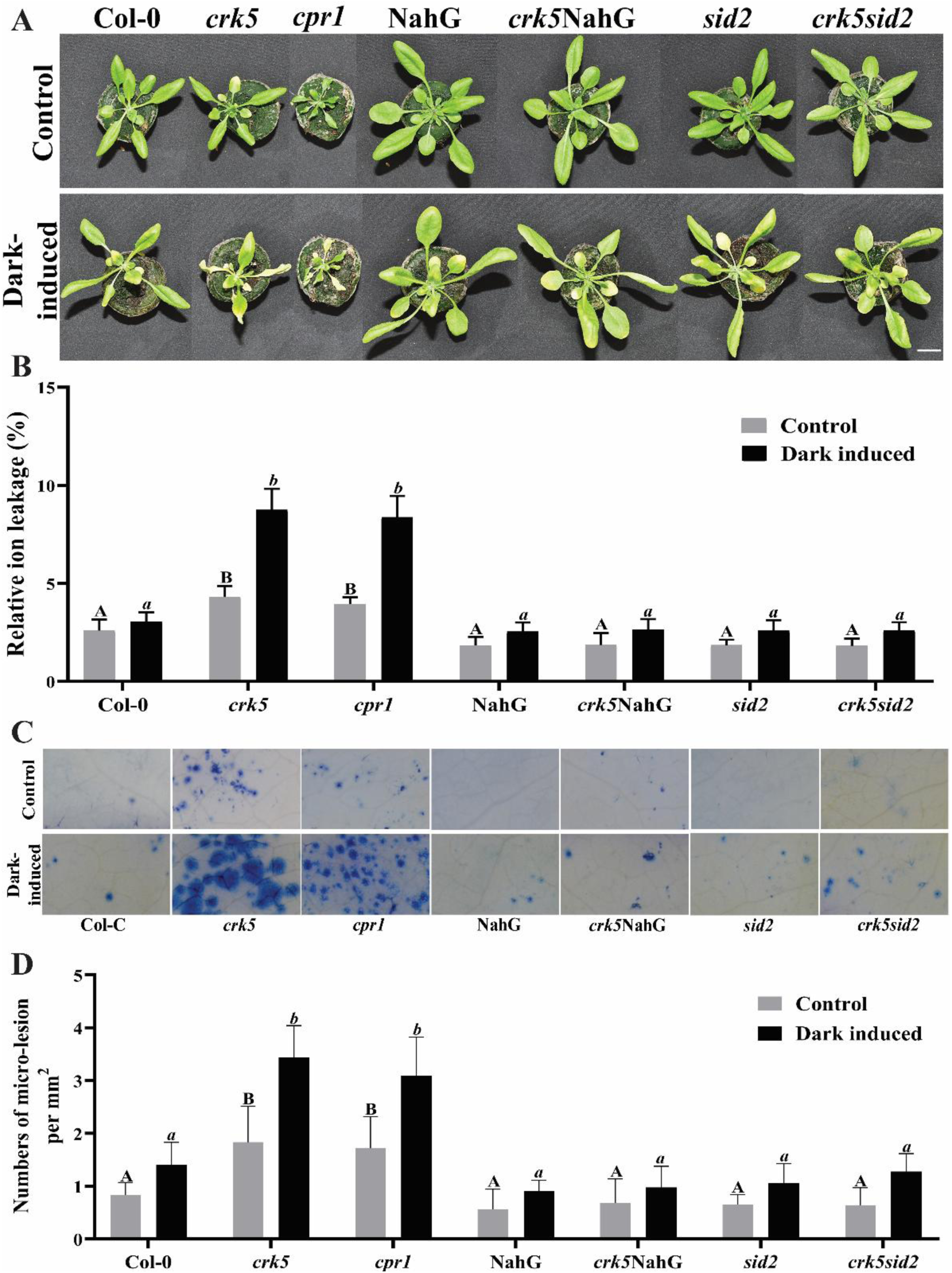
CRK5-dependent blockade of salicylic acid accumulation and signaling during dark-induced senescence and cell death. Morphology (**A**), relative ion leakage (**B**), trypan blue staining for the detection of cell death (**C**), and cell death quantified as micro-lesion number per mm^2^ (**D**) of wild-type, *crk5*, c*pr1*, and salicylic acid-synthesis dysfunction *sid2* mutant or salicylic acid-catabolism transgenic NahG line after exposure for 4 days to permanent darkness. The picture shows a phenotype of 4-week-old plants growing under ambient light (120 μmol m^−2^ s^−1^) conditions and exposed to four days of permanent darkness. White scale bars indicate 1 cm. Mean values (±SD) were derived from 9 plants (n = 9). Statistical analysis was performed using a t-test at a significance level of p < 0.05. Letters A, B, and a, b above the bars indicate homogenous groups, and values sharing common labels (letters) are not significantly different from each other.

### Carotenoid degradation during dark-induced senescence is suppressed by CRK5 and its inhibition of SA-signaling

Considering that SA accumulation under prolonged darkness enhances CD and promotes the aging processes, we examined the effect of these changes on carotene and xanthophyll levels. In our study, we observed a significant reduction in total carotenes (including ɑ-carotenes and ß-carotenes) and xanthophylls (including lutein, violaxanthin, antheraxanthin, and zeaxanthin) in SA-accumulating *crk5* and *cpr1* mutants under both control and dark-induced conditions compared to the wild-type, while introduction of deregulation of SA-synthesis mutant or SA-catabolism transgenic line, respectively (*sid2* and NahG) into *crk5* mutant background reverted its phenotype of accelerated carotenoids and xanthophyll dark-induced degradation to the wild-type phenotype (Fig. 2A, B, and Fig. S2). These results support the role of CRK5 in inhibiting the SA-signaling pathway and its role in protecting SA-dependent degradation of carotenoids and xanthophylls during the onset of DIS.

**Figure 2.**
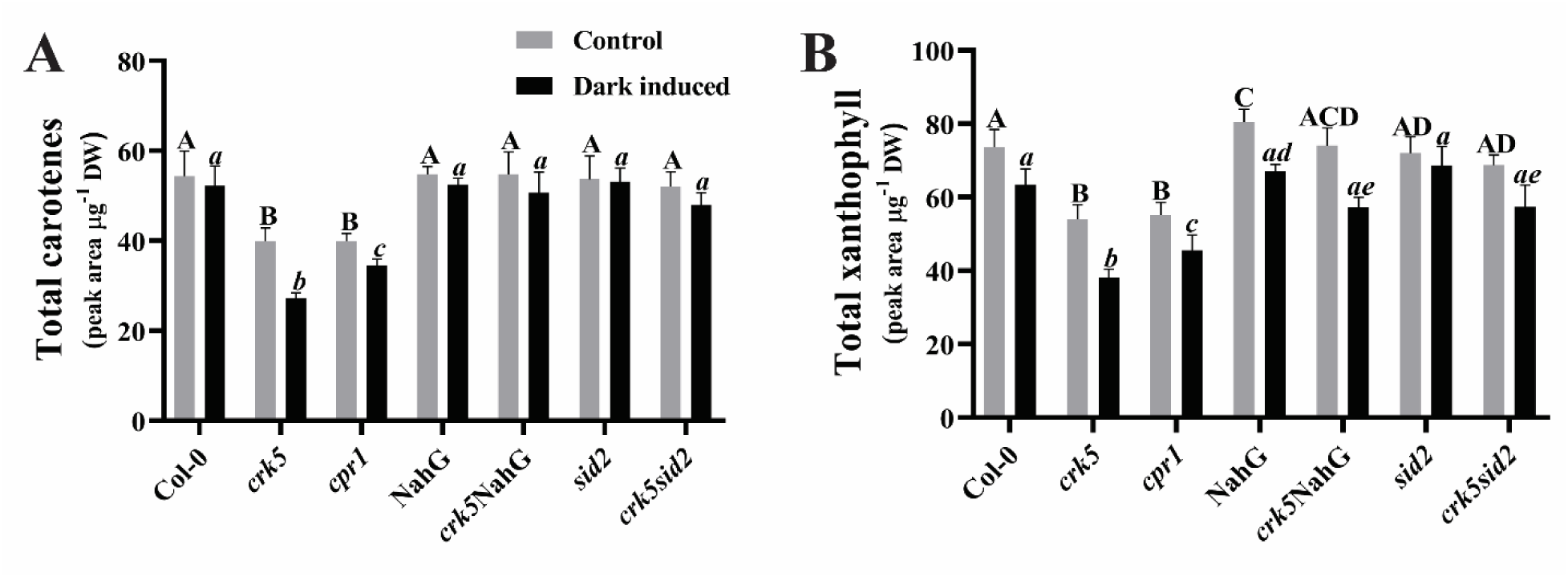
Foliar total carotenes and xanthophyll content of wild-type, *crk5*, *cpr1*, and salicylic acid-synthesis dysfunction *sid2* mutant or salicylic acid-catabolism transgenic NahG line in control conditions and after exposure for 4 days of darkness. Total carotenes content (**A**) and total xanthophyll content (**B**). Mean values (±SD) were derived from 9 plants (n = 9). Statistical analysis was performed using a t-test at a significance level of p < 0.05. Letters A, B, C, D, and a, b, c, d, e above the bars indicate homogenous groups, and values sharing common labels (letters) are not significantly different from each other.

### ROS formation in dark-induced senescence is suppressed by CRK5 and its inhibition of SA-signaling

Since CRK5 was found to affect carotenoid degradation, we further investigated whether it influences the ROS formation and its prevention systems. We found that DPPH and ABTS antioxidant activities were strongly increased in the SA-accumulating *crk5* and *cpr1* mutant lines compared to wild-type, with a further increase under 4 days of DIS. In contrast, SA-synthesis deficient mutant (*sid2*) and transgenic SA-catabolic (NahG) line showed significantly lower antioxidant activity. Introduction of *sid2* and NahG into the *crk5* background reverted antioxidant activity to wild-type phenotype, indicating that CRK5 and its suppression of SA-signaling prevent ROS formation during DIS (Fig. 3A, B). We also observed significantly elevated level of total phenolic compounds in SA-accumulating plants (Fig. 3C). The phenolic profile showed that the levels of phenylpropanoids, flavonols, and anthocyanins in our study were significantly higher in *crk5* and *cpr1*, while in SA-synthesis deficient mutant (*sid2*) and SA-catabolic transgenic (NahG) line, the levels were considerably lower than in wild-type in both control and dark conditions. In double mutant lines, there was reversion in the levels of these phenolic compounds, and the levels were similar to those of SA-synthesis deficient mutant or SA-catabolic line (Fig. 3D–F). These results also indicate a higher level of non-enzymatic antioxidant activity in the SA-accumulating lines during DIS.

**Figure 3.**
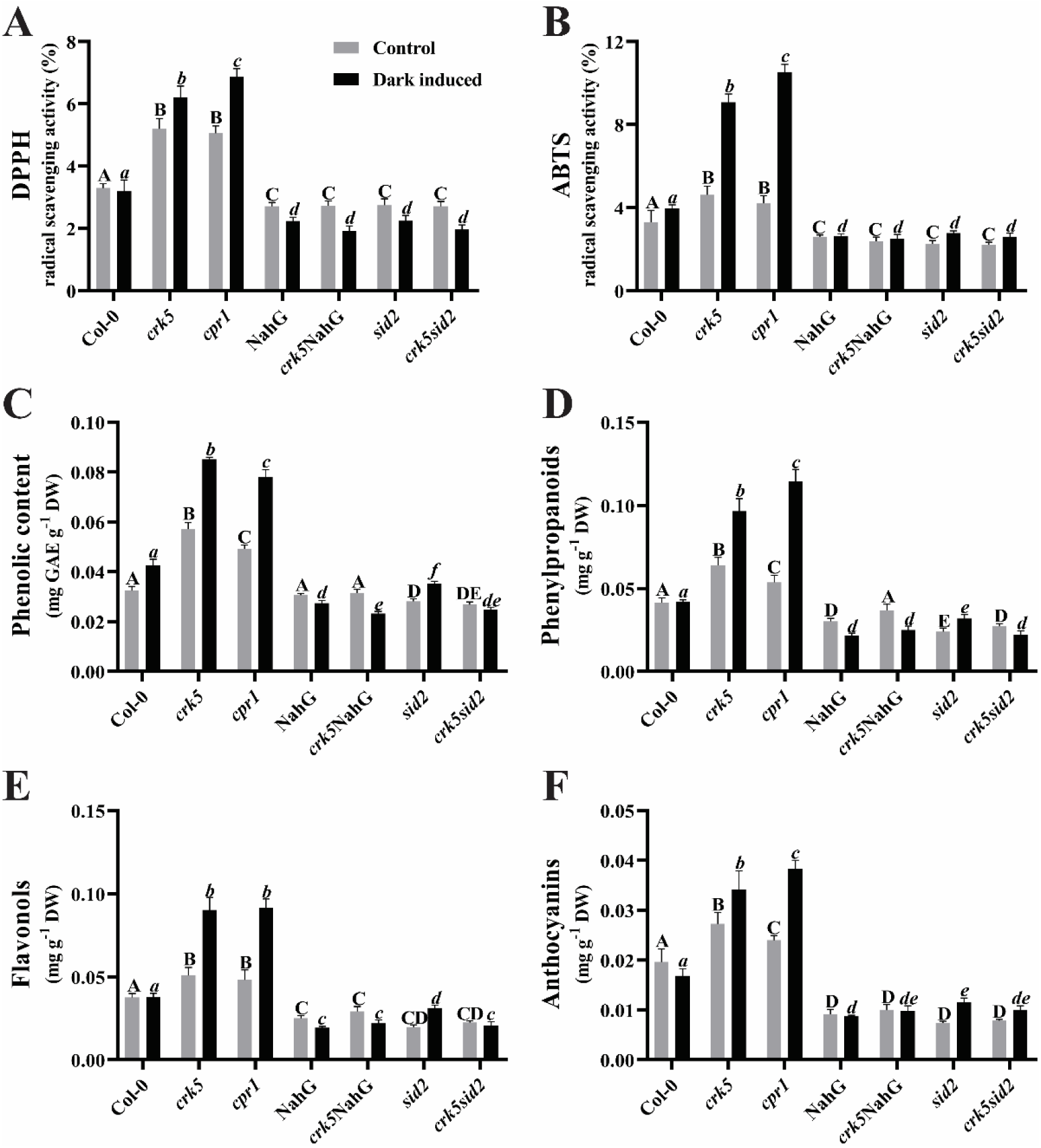
The antioxidant activity of wild-type, *crk5*, *cpr1*, and salicylic acid-synthesis dysfunction *sid2* mutant or salicylic acid-catabolism transgenic NahG line in control and after 4 days exposure to darkness. DPPH (2,2-diphenyl-1-picrylhydrazyl) (**A**), ABTS (2,2′-azinobis (3-ethylbenzothiazoline-6-sulfonic acid)) ROS scavenging activity (**B**), Phenolic (**C**), phenylpropanoids (**D**), flavonols (**E**), and anthocyanins foliar content (**F**). Mean values (±SD) were derived from 9 plants (n = 9). Statistical analysis was performed using a t-test at a significance level of p < 0.05. Letters A, B, C, D, E, and a, b, c, d, e, f above the bars indicate homogenous groups, and values sharing common labels (letters) are not significantly different from each other.

### The ROS level and enzymatic ROS scavengers are dependent on CRK5 and SA-signaling

To assess ROS levels and enzymatic antioxidant system responses to DIS and SA-signaling, we quantified foliar ROS levels and measured the activities of major antioxidant enzymes (APX, CAT, and SOD). Deregulated SA-signaling and -accumulating mutants (*crk5* and *cpr1*) exhibited significantly higher basal and dark-induced H₂O₂ levels than Col-0, whereas SA-synthesis dysfunction mutant (*sid2*) or transgenic SA-catabolic lines (NahG) showed reduced H₂O₂ foliar levels (Fig. 4A, B; Fig. S1). Introduction of *sid2* mutations or NahG activity into the *crk5* background largely suppressed the elevated H₂O₂ levels, demonstrating that CRK5-dependent inhibition of ROS accumulation is mediated by inhibition of SA-signaling. Consistent with ROS patterns, APX, CAT, and SOD enzymatic activities were increased in *crk5* and *cpr1* but remained unchanged in *sid2* and NahG compared with Col-0. These increased activities in *crk5* and *cpr1* were reverted to near wild-type levels in *crk5*NahG and *crk5sid2* double mutants and lines (Fig. 4C–E). Together, these results indicate that CRK5 inhibits the SA-signaling and foliar ROS levels, thus accelerating DIS, characterized by enhanced ROS production and CD.

**Figure 4.**
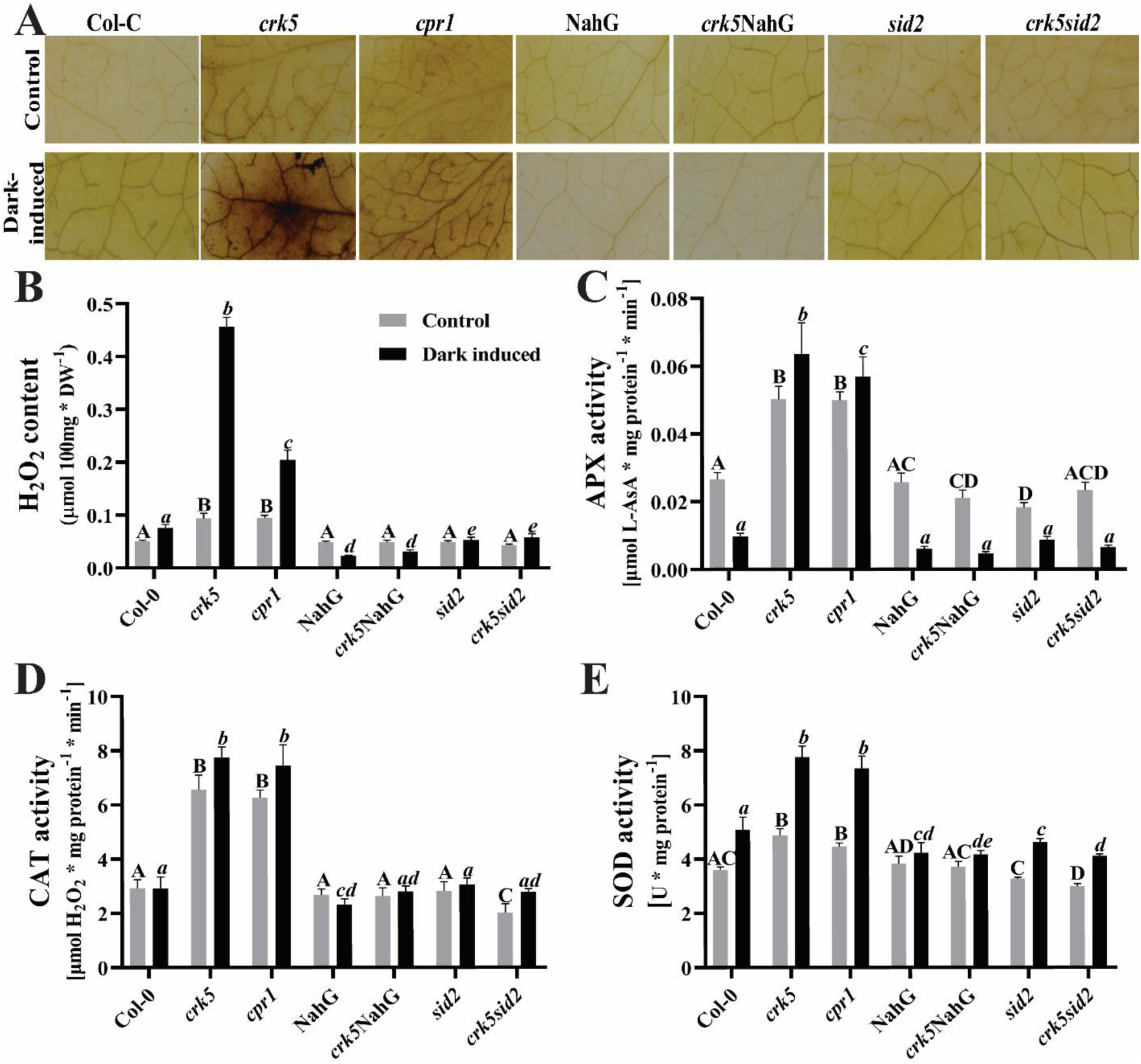
Foliar ROS levels and enzymatic ROS scavenging activity of wild-type, *crk5*, *cpr1*, and salicylic acid-synthesis dysfunction *sid2* mutant or salicylic acid-catabolism transgenic NahG line in control and after exposure for 4 days to darkness. 3,3’diaminobenzidine (DAB) staining showing H_2_O_2_ foliar levels (**A**), H_2_O_2_ foliar content (**B**), APX (**C**), CAT (**D**), and SOD (**E**) foliar enzymatic activity. Mean values (±SD) were derived from 9 plants (n = 9). Statistical analysis was performed using a t-test at a significance level of p < 0.05. Letters A, B, C, D, and a, b, c, d, e above the bars indicate homogenous groups, and values sharing common labels (letters) are not significantly different from each other.

### CRK5 prevents the transcriptomic changes by inhibiting the SA-signaling

To gain a deeper insight into CRK5’s negative regulation of DIS and CD, as well as cellular redox homeostasis, we performed whole-transcriptome sequencing of the analyzed genotypes under ambient light and after four days of continuous darkness. Under control conditions, the SA-synthesis deficient mutant *sid2* and the SA-accumulating mutant *cpr1* exhibited the most pronounced transcriptional changes, with a total of 93 differentially expressed genes (DEGs), underscoring the importance of SA-signaling in senescence- and CD-related processes (Fig. 5A). As expected, senescence- and CD-associated genes showed limited transcriptional regulation under ambient light and photoperiod, with the notable exception of *crk5*, which displayed elevated expression of the senescence markers *SAG12* and *SAG13*, as well as the CD-related genes *MC2* and *MC8*, that is consistent with its accelerated DIS phenotype (Fig. 5C). Gene ontology (GO) enrichment analysis further revealed significant overrepresentation of terms related to senescence, CD, and SA response among genes induced in *crk5* (Fig. S3). Dark-treatment strongly induced senescence- and CD-related transcriptional responses across all genotypes (Fig. 5C). Notably, *crk5* exhibited the most extensive transcriptomic reprogramming, with more than 7,187 DEGs compared with wild-type plants (Fig. 5B). GO analysis of induced genes in *crk5* showed significant enrichment of processes related to pigment catabolism, organ senescence, and responses to light intensity (Fig. S3). Consistent with these findings, *crk5* displayed marked up-deregulation of key senescence regulators (*SAG12*, *SAG29*, *ANAC046*, *NAP*, *WRKY75*, *NYE1*), autophagy-related genes (*ATG5*, *ATG7*, *ATG8a*), and CD-associated genes (*MC2*, *MC8*, *LSD1*, *HRD1A*) (Fig. 5C). In addition, dark-exposed *crk5* plants showed increased expression of genes involved in metabolic and redox-related pathways, including *PALs*, *RBOHs*, *PRXs*, *MPKs*, *GDH2*, *CAT3*, *GPX2*, *GUN1*, *NPR1*, and *CKX2*, suggesting a broad role for SA-signaling in metabolic regulation during accelerated DIS (Fig. 5D). Although the *cpr1* mutant exhibited clear senescence symptoms under darkness, its transcriptional changes were substantially less pronounced than those observed in *crk5*, indicating that CRK5 functions as a major upstream regulator of SA-dependent transcriptional responses during accelerated DIS.

**Figure 5.**
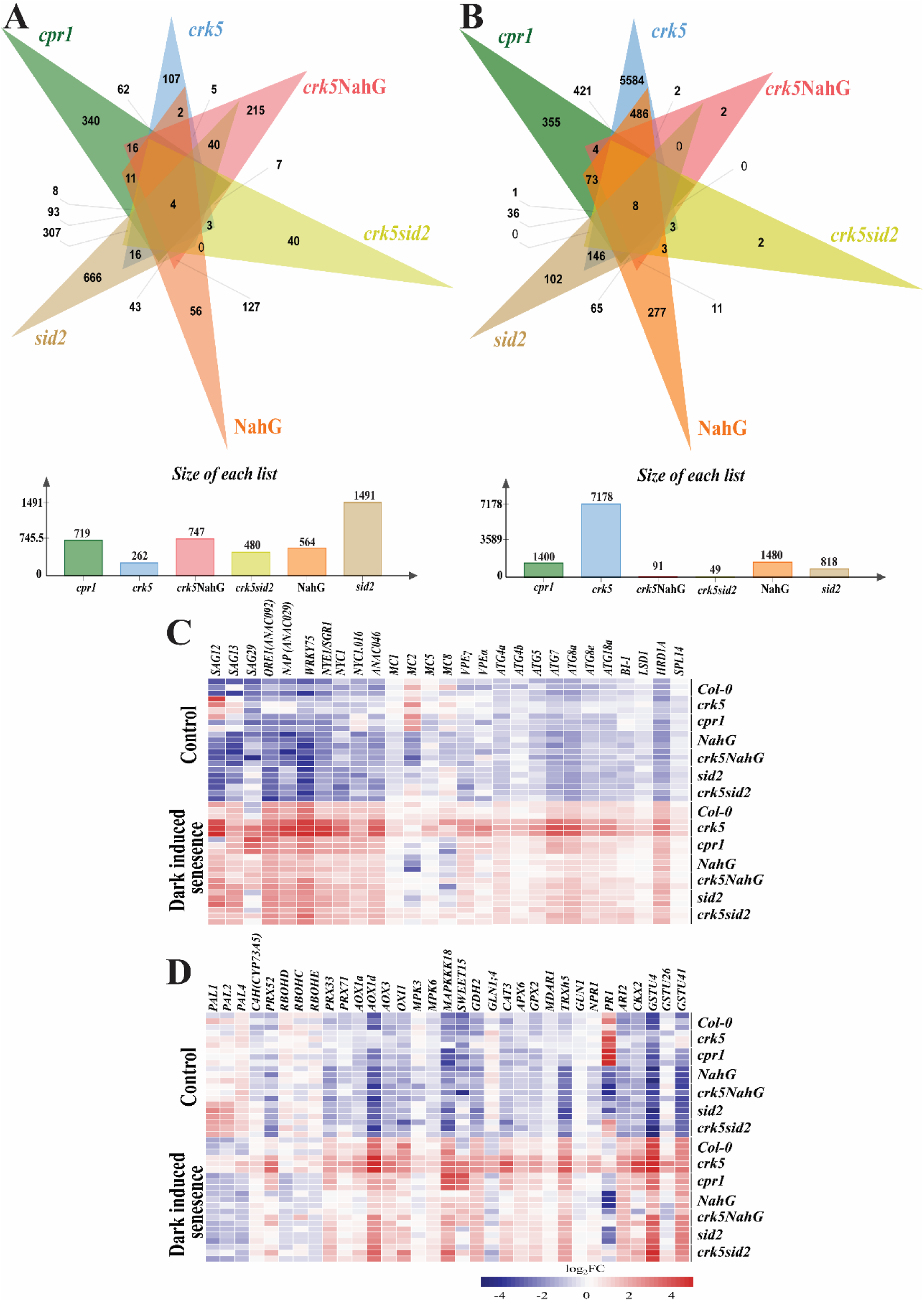
Venn diagram showing commonly and differentially regulated DEGs, and heat maps showing the expression patterns of selected DEGs in wild-type, *crk5*, c*pr1*, and salicylic acid-synthesis dysfunction *sid2* mutant or salicylic acid-catabolism transgenic NahG line under control and after exposure to permanent darkness for 4 days. Number of deregulated transcripts in control (**A**) and number of deregulated transcripts in dark-induced plants (**B**), Selected senescence and CD pathway-related genes (**C**), and metabolic pathways-related genes (**D**).

## Discussion

In the present study, we provided genetic, molecular, and physiological evidence that CRK5 acts as a negative regulator of CD during accelerated DIS through the inhibition of SA-signaling pathway, ROS, and antioxidant homeostasis (Fig. 6). Accelerated senescence and CD in the absence of functional CRK5 in *crk5* mutant was further confirmed by trypan blue staining and quantification of micro-lesions, which revealed a significantly higher number of dead foliar cells (Fig. 1C, D). These observations were consistent with previous studies demonstrating that SA promotes a conditionally dependent hypersensitive disease defense response characterized by spontaneous CD in numerous lesion-mimic mutants [36, 38, 40, 70]. One of our previous studies demonstrated an increased rate of CD in the *lsd1* mutant, characterized by elevated SA levels, under abiotic stress conditions (UV-A+B), further supporting a strong link between SA-signaling and the regulation of CD [68].

**Figure 6.**
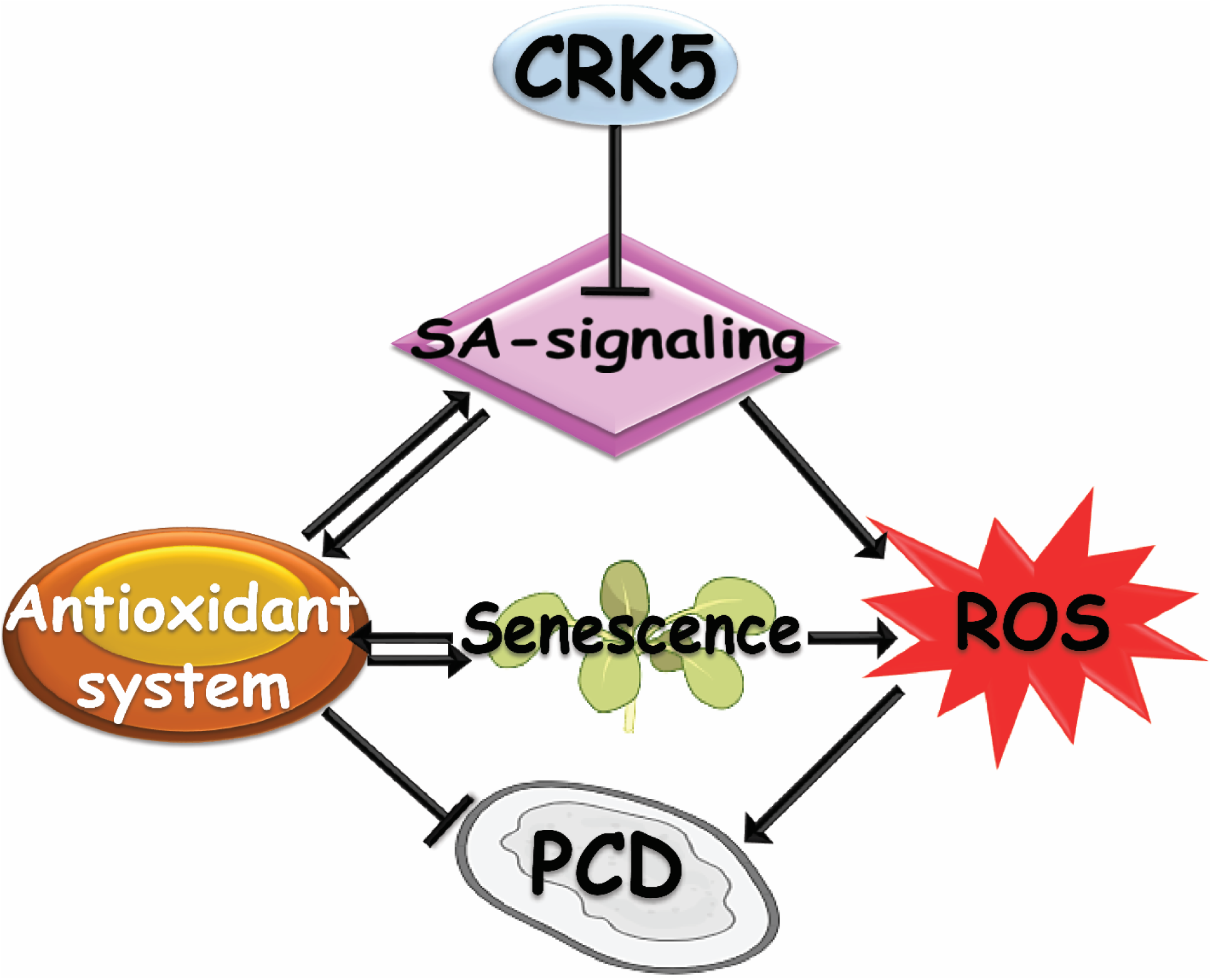
Schematic model illustrating the negative regulatory role of CRK5 in the salicylic acid signaling. The CRK5-driven inhibition of SA-signaling coordinates hormonal, metabolic, and oxidative pathways to maintain leaf viability, providing mechanistic insight into the control of stress-induced senescence and cell death in Arabidopsis.

Previous studies have demonstrated that DIS is accompanied by chlorophyll degradation and major alterations in pigment composition, largely due to impaired photosynthesis and increased photo-oxidative stress. Carotenoids play an essential photo-protective and antioxidant role, and their decline during DIS correlates with enhanced photo-oxidative damage and senescence progression [71, 72]. Consistent with this, we observed a significant reduction in α-and β-carotenes in the SA-accumulating mutants *crk5* and *cpr1* under control conditions, but this reduction was accelerated in dark conditions (Fig. S2A, B). Independent xanthophyll measurements confirmed a similar pattern, with a more pronounced decline during darkness (Fig. 2B and Fig. S2). In contrast, SA-synthesis deficient (*sid2*) and transgenic SA-catabolic (NahG) lines maintained pigment levels comparable to wild-type, and genetic incorporation of these lines into the *crk5* background restored carotenoid content, indicating that CRK5 negatively regulates SA-accumulation and carotenoid depletion during DIS.

SA has been reported to exert concentration- and duration-dependent effects on carotenoid catabolism; while moderate SA levels may stimulate biosynthesis, prolonged or excessive SA-accumulation under stress promotes carotenoid degradation [73, 74]. Accelerated senescence and CD observed in *crk5* and *cpr1* in both conditions likely reflect this imbalance, as carotenoids normally protect against ROS by quenching singlet oxygen, thus reducing free radical formation [19]. Under high SA and oxidative stress, enhanced carotenoid degradation reduces antioxidant capacity, facilitating ROS accumulation and CD progression [75]. SA-mediated regulation of carotenoid cleavage dioxygenases may further link carotenoid turnover to CD signaling through the production of bioactive apo-carotenoids [76].

Consistent with increased oxidative stress, *crk5* and *cpr1* displayed significantly higher DPPH and ABTS antioxidant activities under both control and DIS conditions (Fig. 3A, B). This elevated non-enzymatic antioxidant capacity is characteristic of SA-accumulating and lesion-mimic mutants, where excessive SA induces phenolic biosynthesis as a compensatory response to redox imbalance and CD activation in some cells [28, 38, 68, 70]. Comparable SA-induced enhancements in total antioxidant capacity have been reported under both biotic and abiotic stress conditions, underscoring the dual function of SA as a key signaling molecule and a regulator of cellular redox homeostasis [38, 77]. In agreement with previous reports, *crk5* and *cpr1* accumulated higher levels of total phenolic content and specific subclasses, including phenylpropanoids, flavonols, and anthocyanins, particularly under darkness (Fig. 3C–F). Conversely, SA-synthesis or -catabolic lines showed reduced ROS, CD, and antioxidant capacity, while both parameters were restored to wild-type levels in *crk5sid2* and *crk5*NahG, reinforcing the central role of appropriate SA-signaling in regulating non-enzymatic and enzymatic antioxidant responses during DIS. It is consistent with reports showing that suppression of SA-biosynthesis or -signaling leads to decreased antioxidant potential and reduced CD [40, 78]. Furthermore, the restoration of wild-type antioxidant capacity in *crk5sid2* and *crk5*NahG double mutants supports the idea that the enhanced antioxidant activity in *crk5* plants is due to deregulation of cellular ROS levels as a secondary effect of SA-dependent signaling and CD activation.

SA is also known to modulate enzymatic antioxidant defenses. Exogenous SA application is related to a significant upregulation of major enzymatic antioxidants, including SOD, APX, and CAT [79–81]. Consistent with this, we observed a significantly higher H_2_O_2_ level in SA-accumulating mutants and reduced ROS formation in SA-deficient mutants (Fig. 4A and B). The progression of DIS is naturally linked to the loss of cellular redox control, whereas the rapid accumulation of ROS serves as a critical signaling mediator for CD. At lower, protective physiological concentrations, exogenous SA application has been shown to counteract senescence promoting signals, notably by strengthening the overall antioxidant defense system. This protective effect involves enhancing the activity of key scavenging enzymes such as APX, CAT, and SOD, thereby reducing deleterious ROS accumulation [6, 14, 27]. In our study, we also found an elevated activity of antioxidant enzymes APX, CAT, and SOD in SA-accumulating lines (Fig. 4C–E). This upregulation likely represents a compensatory response to excessive ROS accumulation rather than a protective suppression of senescence.

Our transcriptomic analyses further position CRK5 as a pivotal upstream negative regulator of SA-dependent transcriptional reprogramming during DIS. consistent with previous reports showing that SA contributes to developmental and stress-induced leaf aging [24, 82, 83]. Under ambient conditions, SA-synthesis deficient (*sid2*) and SA-accumulating (*cpr1*) mutants exhibited the most pronounced transcriptional divergence, highlighting the importance of SA-signaling in CD and antioxidant regulation. Notably, *crk5* and *cpr1* showed accelerated induction of senescence markers (*SAG12*, *SAG13*) and CD-related genes (*MC2, MC8*), consistent with their elevated SA levels. Upon dark treatment, the *crk5* plants displayed the most extensive deregulated transcriptomic remodeling, with over 7,100 DEGs, including strong induction of senescence-, autophagy-, and CD-associated genes (*SAGs*, *ANACs*, *MCs*, *VPEs*, *ATGs*, *LSD1*) (Fig. 5C), which is consistent with previous studies linking CD progression to nutrient remobilization, active catabolic remodeling and senescence [84–86]. At the same time, genes involved in redox regulation and antioxidant catabolism (*PALs*, *PRXs*, *AOXs*, *CAT3*, *APX6*, *GPX2*) were strongly induced, indicating tight coupling between the onset of CD and antioxidant homeostasis.

The GO enrichment analysis corroborated these findings, revealing significant overrepresentation of senescence, ROS response, detoxification, metabolic catabolism, and light-response pathways among *crk5*-induced genes (Figs. 5B, S3), consistent with global transcriptional reprogramming during leaf aging [87]. The comparatively modest transcriptional response of *cpr1* under darkness, despite clear senescence symptoms, suggests that CRK5 functions upstream of CPR1 in orchestrating SA-driven inhibition of CD and redox regulation. Importantly, the near-complete reversion of *crk5* phenotypes and transcriptomic changes in SA-synthesis or -catabolism deficient double mutants confirms that CRK5-mediated inhibitory effects on CD and antioxidant homeostasis are largely SA-signaling dependent (Fig. 6).

## Conclusions

In summary, our findings identify CRK5 as a central inhibitor of SA-signaling-dependent CD and antioxidant redox homeostasis during DIS. Loss of CRK5 leads to enhanced SA-accumulation, accelerated CD, and destabilization of carotenoid pools, phenolic metabolism, and enzymatic antioxidant balance, thereby exacerbating oxidative stress and senescence progression. Transcriptomic analyses further position CRK5 upstream of SA-driven CD signaling, coordinating antioxidant defense and catabolic reprogramming during senescence. Collectively, our findings establish CRK5 as a key molecular hub where hormonal signaling, redox regulation, and stress-induced CD integrate to maintain leaf metabolic stability.

## Declarations

### Ethics approval and consent to participate

Not applicable.

### Consent for publication

Not applicable.

### Availability of data and materials

The raw RNA-seq (fastq) data are deposited in the NCBI database (BioProject: PRJNA1389866).

### Competing interests

The authors declare no competing interests.

### Funding

This research was funded by the Polish National Science Centre (Opus 20 UMO 2020/39/B/NZ3/02103), granted to S.K.

### Authors’ Contributions

Conceptualization and proposed working hypothesis SK and MK; experimental design of the research SK, PB, and MK; resources, performed the research, and RNA-seq data analysis MK, AR, and RZG; Supervision SK and PB; writing the original draft MK and PB; revision of the manuscript SK.

## Acknowledgements

Not applicable.

## Notes

### Competing Interest Statement

The authors have declared no competing interest.

