## Supplementary file for "CRK5 preserves antioxidant homeostasis and prevents cell death during dark-induced senescence through inhibiting the salicylic acid signaling pathway"

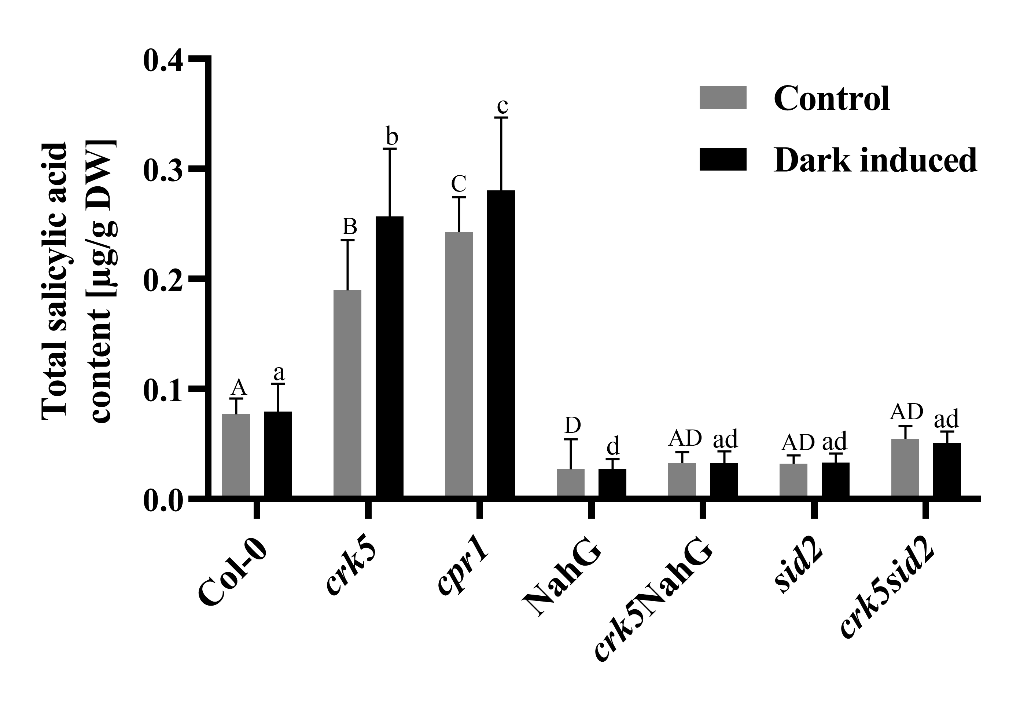

**Figure S1.** Total salicylic acid (SA) content of the wild-type, *crk5*, c*pr1*, and salicylic acid-synthesis dysfunction *sid2* mutant or salicylic acid-catabolism transgenic NahG line in control and after exposure for 4 days to permanent darkness. Mean values (±SD) were derived from 9 plants (*n* = 9). Statistical analysis was performed according to a *t*-test at a level of *p* < 0.05. Letters A, B, C, D, and a, b, c, d above the bars indicate homogenous groups, and values sharing common labels (letters) are not significantly different from each other (Modified from Kamran et al, 2025, accepted in Plant Physiology).

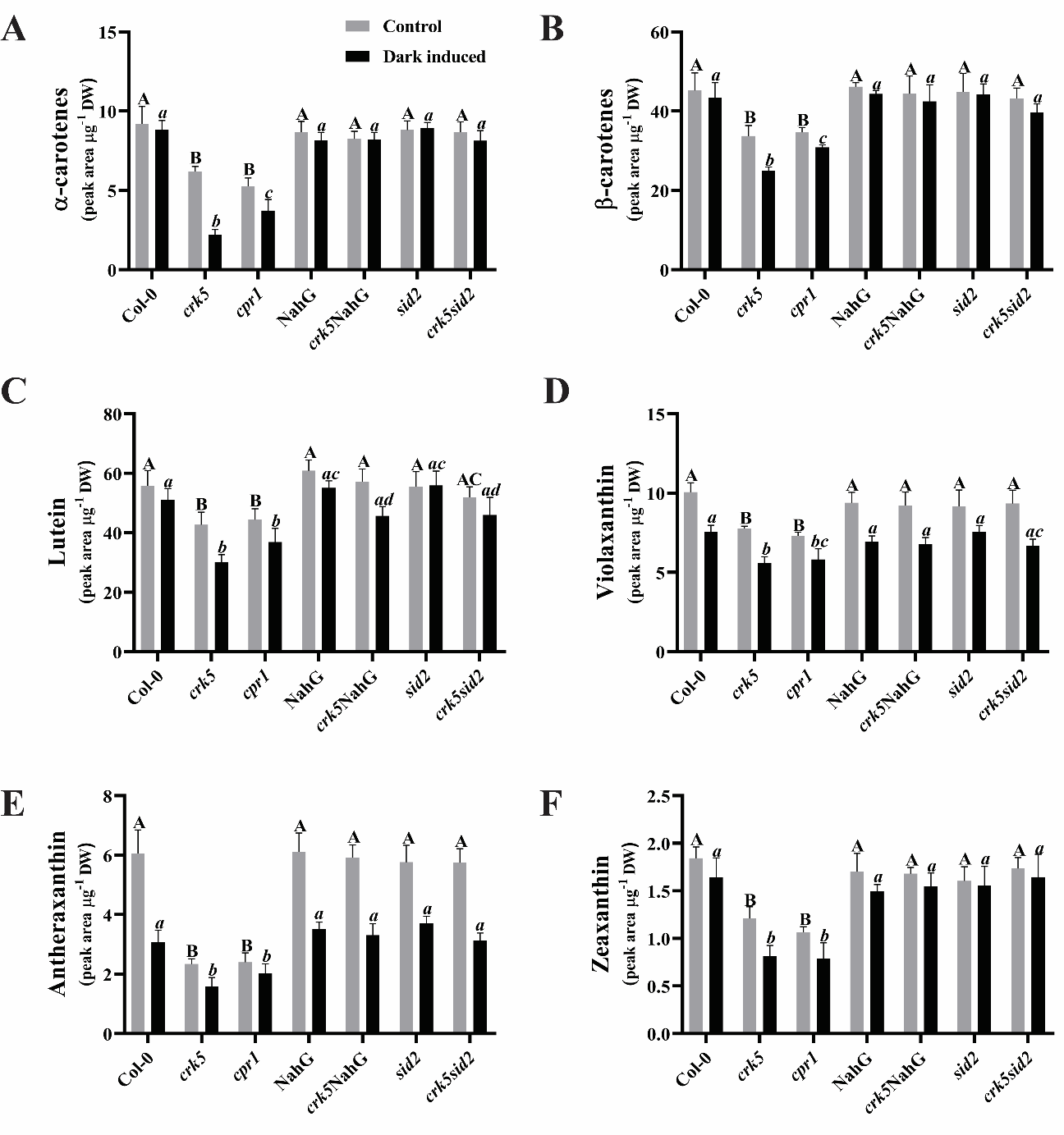

**Figure 2.** Foliar carotenes and xanthophylls content of wild-type, *crk5*, c*pr1*, and salicylic acid-synthesis dysfunction *sid2* mutant or salicylic acid-catabolism transgenic NahG line in control and after exposure for 4 days to permanent darkness. ɑ-carotenes (**A**), ß-carotenes (**B**), lutein (**C**), violaxanthin (**D**), antheraxanthin (**E**), and zeaxanthin (**F**). Mean values (±SD) were derived from 9 plants (n = 9). Statistical analysis was performed using a t-test at a significance level of p < 0.05. Letters A, B, C, and a, b, c, d, above the bars indicate homogenous groups, and values sharing common labels (letters) are not significantly different from each other.

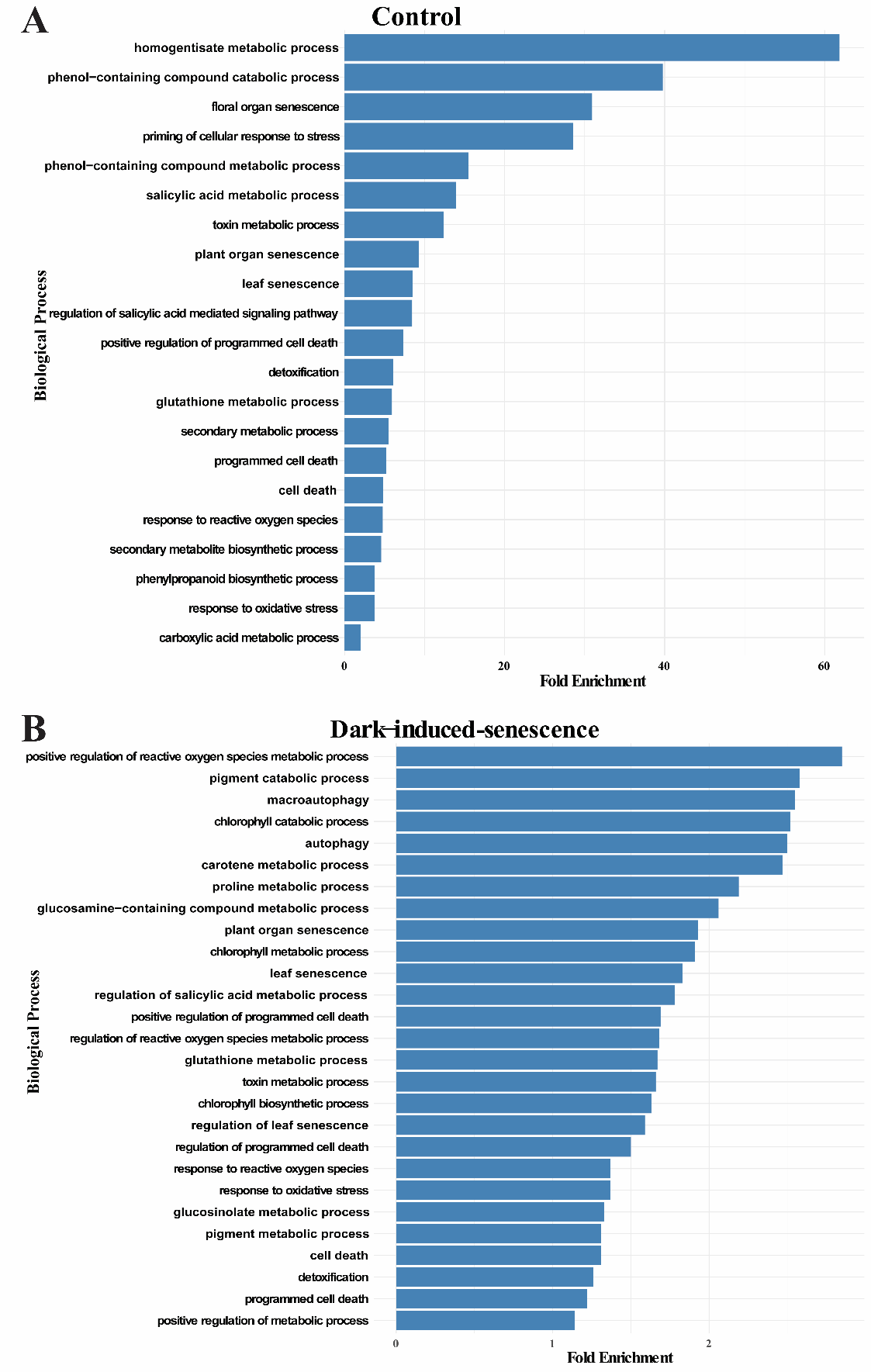

**Figure S3.** Gene ontology (GO) analysis of genes significantly enriched in *crk5* compared to Col-0 in the RNA-seq experiment in control (**A**) and dark-induced (**B**) conditions.

**Table S1.** Primers used in this study.

| **Gene** | **AGI Code** | **Sequence LP Primer** | **Sequence RP Primer** |
| --- | --- | --- | --- |
| *CRK5* | AT4G23130 | AGGAGATCTCTCGCCAGAATC | CGATAGTCTCTTCACGGCAAC |
| NahG | Transgene line (M60055) | ACTCTGCCGCTACTCCCATA | CGAGCCCTAGGTACATCTGC |
| *SID2* | AT1G74710 | AATTGGCAGGGAGACTTACGAA | TCCTCTATCGAATGATTCTATCTCCTT |

**Table S2.** *Accession Numbers* for Fig. 5C and D.

| **Senescence Marker Genes** | **AGI Code** | **SA Signaling Pathway Genes** | **AGI Code** |
| --- | --- | --- | --- |
| *SAG12* | AT5G45890 | *AAO3* | AT2G27150 |
| *SAG13* | AT2G29350 | *AOX1a* | AT3G22370 |
| *SAG29* | AT5G13170 | *AOX1d* | AT1G32350 |
| *ORE1 (ANAC092)* | AT5G39610 | *APX6* | AT4G32320 |
| *NAP (ANAC029)* | AT1G69490 | *ARR2* | AT4G16110 |
| *WRKY75* | AT5G13080 | *C4H (CYP73A5)* | AT2G30490 |
| *NYE1/SGR1* | AT4G22920 | *CAT3* | AT1G20630 |
| *NYC1* | AT4G13250 | *CKX2* | AT2G19500 |
| *ANAC016* | AT1G34180 | *DAO1* | AT1G14100 |
| *ANAC046* | AT3G04060 | *GDH2* | AT5G07440 |
| *MC1* | AT1G02170 | *GLN1.4* | AT5G16570 |
| *MC2* | AT4G25110 | *GPX2* | AT2G31570 |
| *MC5* | AT5G64240 | *GSTU26* | AT1G17180 |
| *MC8* | AT1G16420 | *GSTU4* | AT2G29470 |
| *VPEγ* | AT4G32940 | *GSTU41* | AT2G29460 |
| *VPEα* | AT2G25940 | *GUN1* | AT2G31400 |
| *VPEδ* | AT3G20210 | *MAPKKK18* | AT1G05100 |
| *ATG4a* | AT2G44140 | *MDAR1* | AT3G52880 |
| *ATG4b* | AT3G59950 | *MPK3* | AT3G45640 |
| *ATG5* | AT5G17290 | *MPK6* | AT2G43790 |
| *ATG7* | AT5G45900 | *NPR1* | AT1G64280 |
| *ATG8a* | AT4G21980 | *OXI1* | AT3G25250 |
| *ATG8e* | AT4G16520 | *PAL1* | AT2G37040 |
| *ATG18a* | AT3G62770 | *PAL2* | AT3G53260 |
| *BI-1* | AT5G47120 | *PAL4* | AT3G10340 |
| *LSD1* | AT4G20380 | *PR1* | AT2G14610 |
| *HRD1A* | AT1G18270 | *PRX33* | AT3G49120 |
| *SPL14* | AT1G20980 | *PRX52* | AT5G05340 |
|  |  | *PRX53* | AT5G05340 |
|  |  | *PRX71* | AT5G64100 |
|  |  | *RBOHC* | AT5G51060 |
|  |  | *RBOHD* | AT5G47910 |
|  |  | *RBOHE* | AT1G19230 |
|  |  | *SWEET15* | AT5G13170 |
|  |  | *TRXh5* | AT1G45145 |
